# Super resolution ultrasound imaging Using deep learning based micro-bubbles localization

**DOI:** 10.1101/2022.09.21.508222

**Authors:** Feixiao Long, Weiguang Zhang

## Abstract

Super resolution ultrasound imaging has shown its potential to detect minor structures of tissues beyond the limit of diffraction and achieve sub-wavelength resolution through localizing and tracking the ultrasound contrast agents, such as micro-bubbles. Normally, one important step of super resolution ultrasound imaging, micro-bubbles localization is implemented through conventional computer vision techniques, such as local maxima detection etc. However, these classical techniques are generally time consuming and need fine-tuning multiple parameters to achieve the optimal results. Hence, in the manuscript, a deep learning based micro-bubbles localization is proposed, trying to replace or simplify the complex operations of classical methods. The efficiency of our proposed models is preliminarily proved through 2022 ultra-SR challenge.

## I. Introduction

Super resolution (SR) ultrasound imaging has been proved to be a valuable ultrasound modality to detect fine and detailed structure of blood vessel or small animal organ beyond the limit of sound wave diffraction [1]. Currently, SR ultrasound imaging has been applied to differentiate malignant and benign tissues in rats [2] or human breast [3]. It is also applied to neurology to detect small vessels pathology, which can be viewed as an indicator for brain dysfunction [4].

The manuscript is a brief summary of results submitted to 2022 ultra-SR challenge (https://ultra-sr.com/, abbreviated as challenge in the following parts) ^1^. Normally, the SR images are formed through localizing and tracking some ultrasound contrast agents, such as micro-bubbles (MBs). The MBs localization is conventionally implemented through local maxima detection [6], 2-D Gaussian fitting after deconvolution with point spread function (PSF) [7] etc. Some MBs localization methods through deep learning are recently proposed [8], [9]. Deep learning based technique is also the main topic and method of our manuscript. To the tracking part, Monte Carlo based on Markov chain [10] and maximum intensity crosscorrelation between frames [11] etc. are proposed to solve the issue. Besides, Hungarian algorithm [6] based approach is also employed to track MBs.

The main contributions of our proposed method are (1) more efficient and powerful variants of conventional U-net, which, to our best knowledge, were not employed in MBs localization previously are tailored to solve the issue; (2) Dice loss is added to the total loss function for better convergence. Besides, through adjusting the threshold for segmentation during training (see Sec. II.A), accuracy of pinpointing the local maxima position can be easily controlled, hence increasing the overall model performance.

## II. Methods

Common steps of SR ultrasound imaging includes B images generation, slow time filtering for separation of MBs and tissue background, MBs localization, MBs tracking and tracks accumulation [1] as well as necessary motion compensation between frames. Below we mainly state the details of MBs localization and tracking (Fig. 1).

**Fig. 1.**
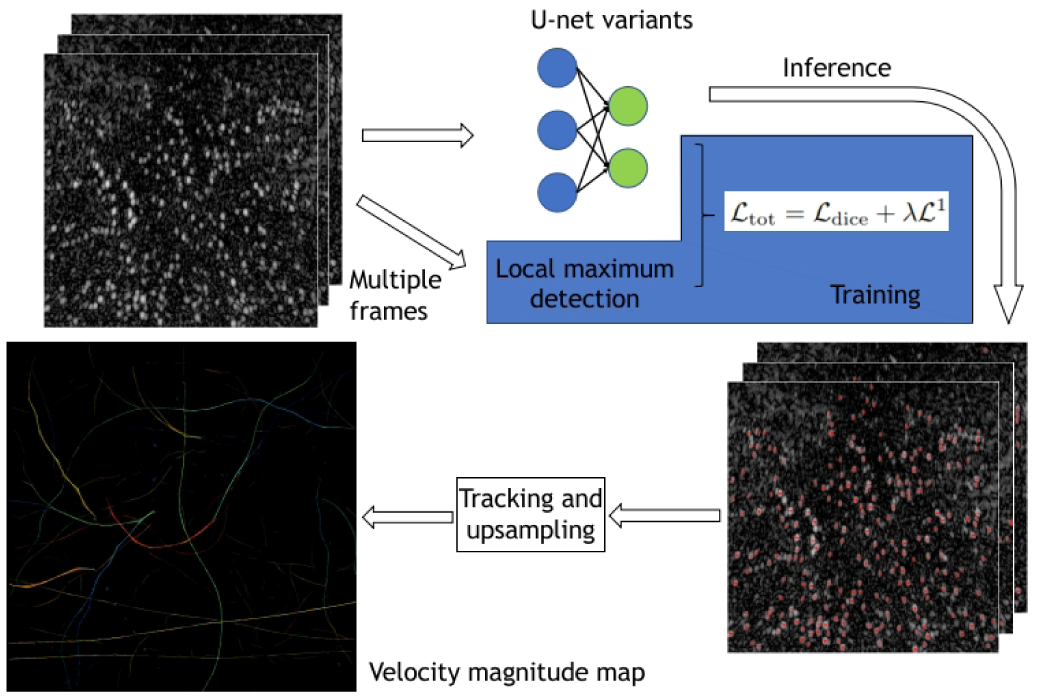
SR ultrasound imaging processing pipeline in our methods.

### A. Deep learning based localization

As illustrated above, a deep-learning based localization method was proposed to replace conventional time consuming computer vision (CV) operations, such as blob or local maxima detection etc. In our method, the MBs localization was transferred to heatmap local maxima detection hence it is nature and straightforward to use deep learning based technique.

## Training data generation

First we would like to point out unlike methods shown in [12], training data generated by simulations were not employed in our experiments for following consideration. It seems that oscillations of MBs are not considered in above mentioned paper and essentially, MBs and normal scatters suffer different underlying physical disciplines when insonified by ultrasound beam [13]. Therefore, we believe accurate simulations with oscillations of MBs needs to be investigated in our future work however it is beyond the scope of this manuscript. Hence, inspired by the work of Deep-stochastic optical reconstruction microscopy (Deep-STORM) [14], we decided to directly use real ultrasound images with patches cropping mode. Our mode can greatly increase the number of training samples with appropriate data augmentations, such as contrast adjust and image smoothing etc. More important, our method does not induce any simulation error.

Since in the challenge, only 4 datasets (two synthetic and two in-vivo datasets) were provided and their appearance are quite different (see Fig. 2), herein in our solution, 4 models were separately trained. To synthetic dataset, first 20 frames from total 500 frames were selected to generate training patches. To in-vivo rat brain dataset, randomly selected 10 videos (800 frames total) from 100 videos (8,000 frames) were employed to train the corresponding model. For in-vivo human lymph node dataset, first 594 frames from video 1 (total 3 videos with 1,382 frames) were used. Because no ground truth of locations of MBs were provided in the challenge, local maxima detection was employed with the help of Matlab buildin function imregionalmax.m [6].

**Fig. 2.**
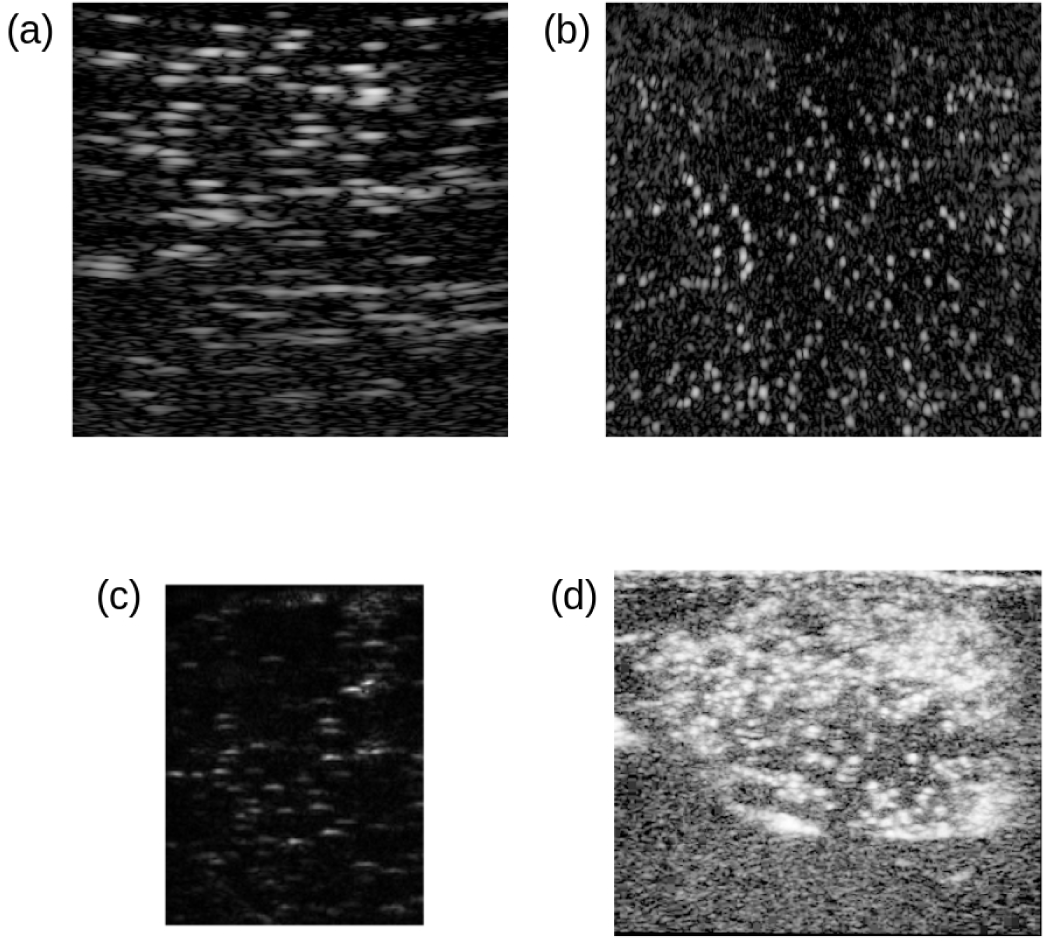
Video frames from the challenge dataset. (a) Frame 1 of synthetic dataset 1. (b) Frame 1 of synthetic dataset 2. (c) Frame 1 of in-vivo rat brain dataset video 1. (d) Frame 300 of in-vivo lymph node dataset video 1.

After getting the positions of MBs, Gaussian function with kernel size 3 *×* 3 and *σ* = 1 was convolved with the localization images, generating the heatmaps which were employed as the labels of training samples. In order to further increase the accuracy of MBs localization, all the images were up-sampled to 3 times larger. Besides, the images were interpolated to the same pixel size in axial and lateral direction with bi-linear method before cropping for the dataset with anisotropic pixel size, such as synthetic dataset 1. Note that the ratio between training samples and total samples is less than normal deep learning training strategy (normally 70% or more) due to similar appearance of individual MB.

### Model architecture selection

Because the MBs localization has been transferred to heatmap local maxima detection, widely-employed encoder-decoder structure, e.g. Unet [15], was employed as our baseline model. Right now, popular transformer-based models are not considered in this challenge due to their requirements of large amount of training samples in general, but will be investigated in our future work. Two variants of classical U-net were mainly explored in our solutions, nnU-net [16] and U-net with ImageNet pretrained ResNet-18 as encoder backbone. nnU-net adopts self-adapting framework to dynamically trade off the batch size and the model capacity, achieving better performance than conventional U-net. For U-net with pretrained ResNet18, spatial and channel attention sub-module [17] was applied in decoder branch, automatically adjusting the spatial pixels and channel contributions to final predictions through trainable weights.

### Loss function

Besides conventional *L*^1^ loss [12], dice loss was added to the total loss, increasing the convergence speed of training in our experiments. When using dice loss, one threshold was pre-set to segment the heatmap (label) to foreground and background. Herein, setting higher value of threshold will ideally increase the accuracy of localization. The total loss used in the challenge was

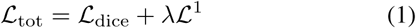

In the equation, *λ* was empirically set to 2.0 and the normal dice loss [18] and *L*^1^ loss were employed.

### Training and inference details

The samples generated in Sec. II.A.1) were split into training subset and validation subset with proportion 8 : 2. The model with minimum validation loss was saved and used for predictions. During the training phase, widely used ADAM optimizer was employed using PyTorch 1.11. The models were implemented with the help of MONAI 0.8.1 [19] and Segmentation models 0.3.0 [20]. All the training and inference were implemented on one Nvidia GPU (GeForce, RTX 2060).

Before predictions, the interpolated test images (with isotropic pixel size) were also up-sampled to 3 times larger. The sliding window (256 *×* 256) fashion with window overlapping ratio 0.25 over up-sampled images was performed after that.

### B. Linear sum assignment based tracking

In the tracking part, conventional tracking algorithm through linear sum assignment (LSA) was employed because of the following considerations. The appearance of MBs are quite similar (see Fig. 2) and identification of single MB only through the image features, whatever manually selected or learned by deep learning, is theoretical difficult. Therefore using LSA is more straightforward and nature although it may be time consuming [6]. The distances between potential localized MBs from previous step and current step were used as the entries of cost matrix and conventional Hungarian algorithm was employed to search the global minimum to the cost function of LSA. After the tracking paths were completed, the MBs density and velocity images could be calculated accordingly.

## III. Results

### A. Accuracy of localization

Fig. 3 to Fig. 4 show the estimated MBs locations with two neural nets for different dataset respectively. For synthetic dataset 1 (see Fig. 3), it seems that U-net with ResNet-18 as backbone predicts the MBs locations with lower intensity more than nnU-net (Fig. 3 (a)). Besides, most predicted positions of MBs by two nets correspond whatever for synthetic dataset 1 or dataset 2.

**Fig. 3.**
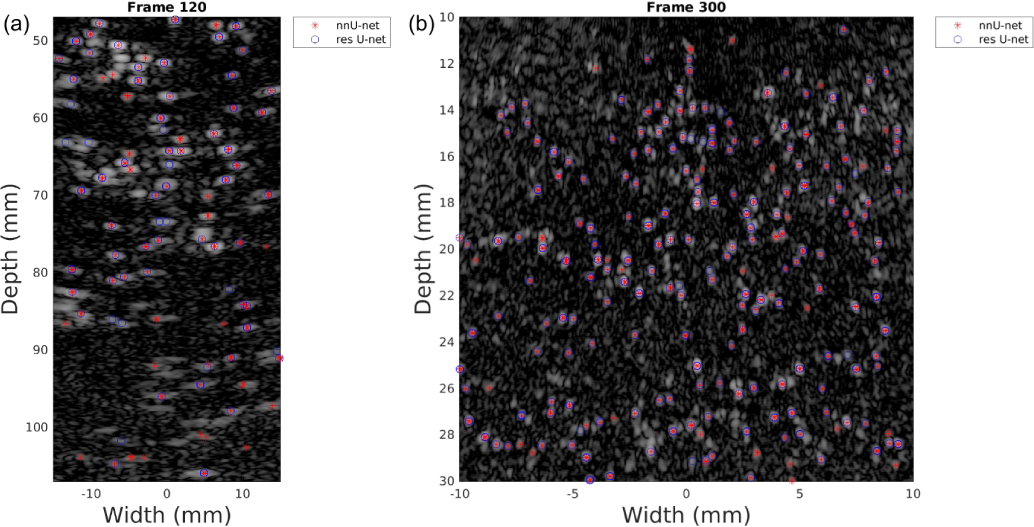
Localization results for synthetic dataset. (a) Synthetic dataset 1. (b) Synthetic dataset 2.

**Fig. 4.**
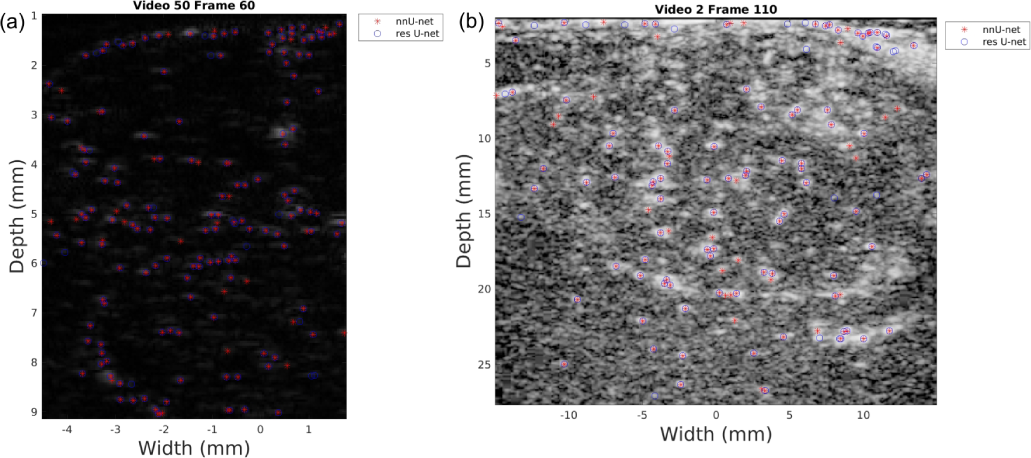
Localization results for in-vivo dataset. (a) In vivo rat brain. (b) In vivo lymph node.

To in-vivo rat brain data, visually most bright points which may correspond the MB locations are detected. To in-vivo lymph node, since the dataset was log compressed, it looks like more noisy. However, at least from Fig. 4, the detection results still seem reasonable for two nets.

### B. Super resolution images

Considering the large size of super resolution images (10 times larger than original size), only portions of images are show in Fig. 5 and Fig. 6. Fig. 5 and Fig. 6 show the MBs density and velocity magnitude map in in-vivo rat brain cortex parts, respectively. It is obvious that some minor vessel structures can be clearly seen in the figures. Since obvious minor structure cannot be seen in portion of lymph node images, the images are rescaled to 0.3 of original ones to show the complete view in Fig. 7 and Fig. 8. From the figures, it can be seen that in the middle part of images, minor bright spots which may represent small lymph nodes can be easily seen. However, we don’t know the actual structure of whole lymph node and this dataset is the only log compressed one in the challenge (also more noisy than other datasets visually), some parameters based on prior information, especially tracking related parameters such as minimum length of single track etc., may not be set to the optimal values.

**Fig. 5.**
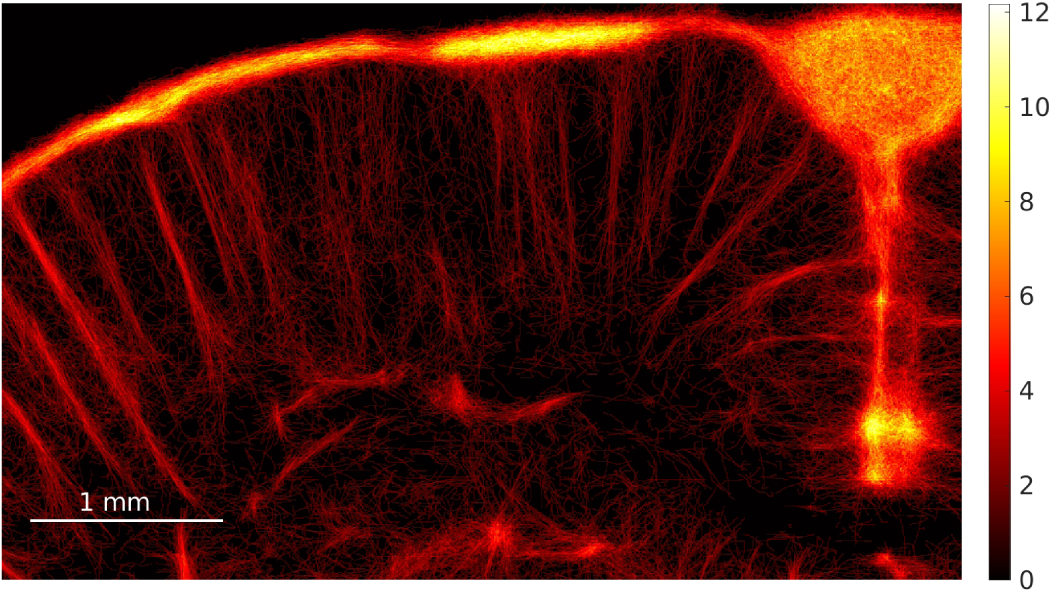
Rat brain MBs density map (a.u.).

**Fig. 6.**
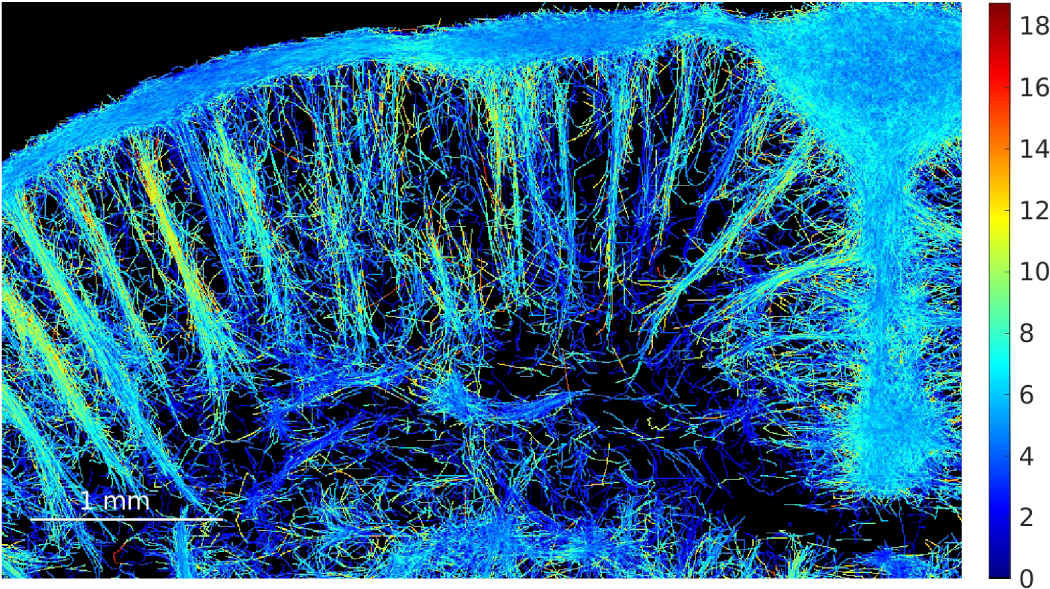
Rat brain MBs velocity magnitude map (mm/s).

**Fig. 7.**
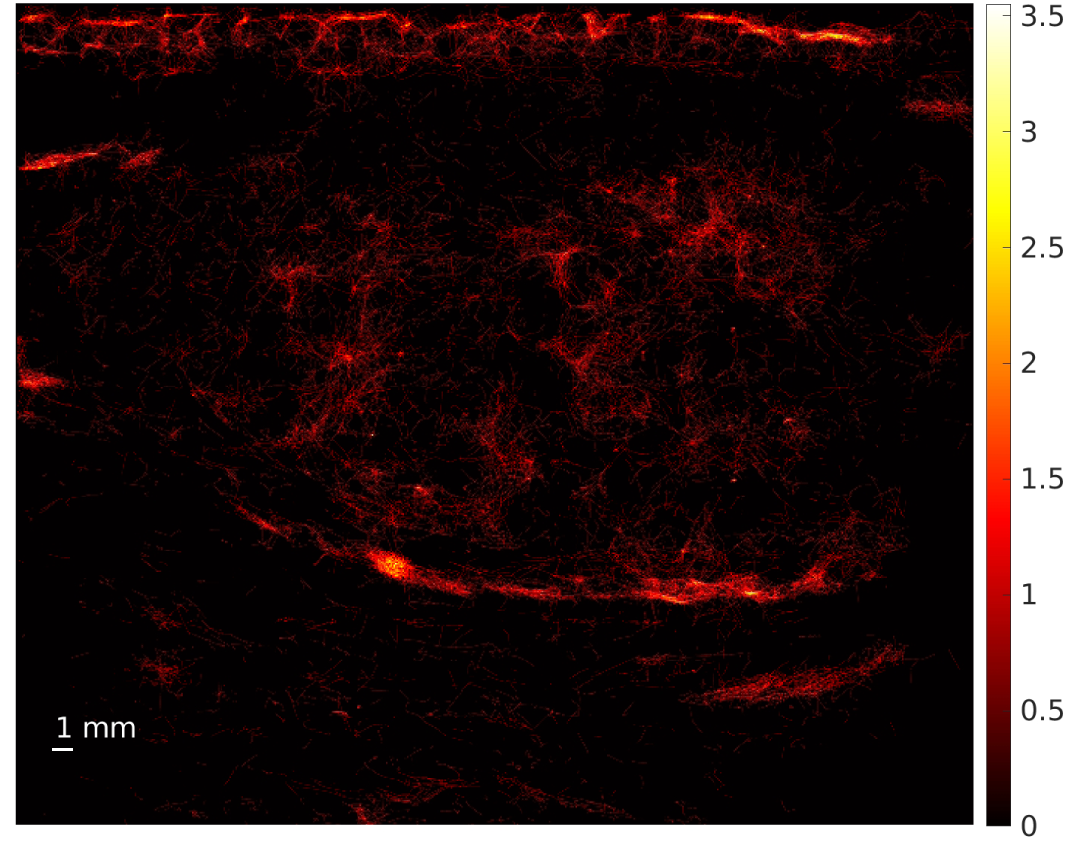
Lymph node MBs density map (a.u.).

**Fig. 8.**
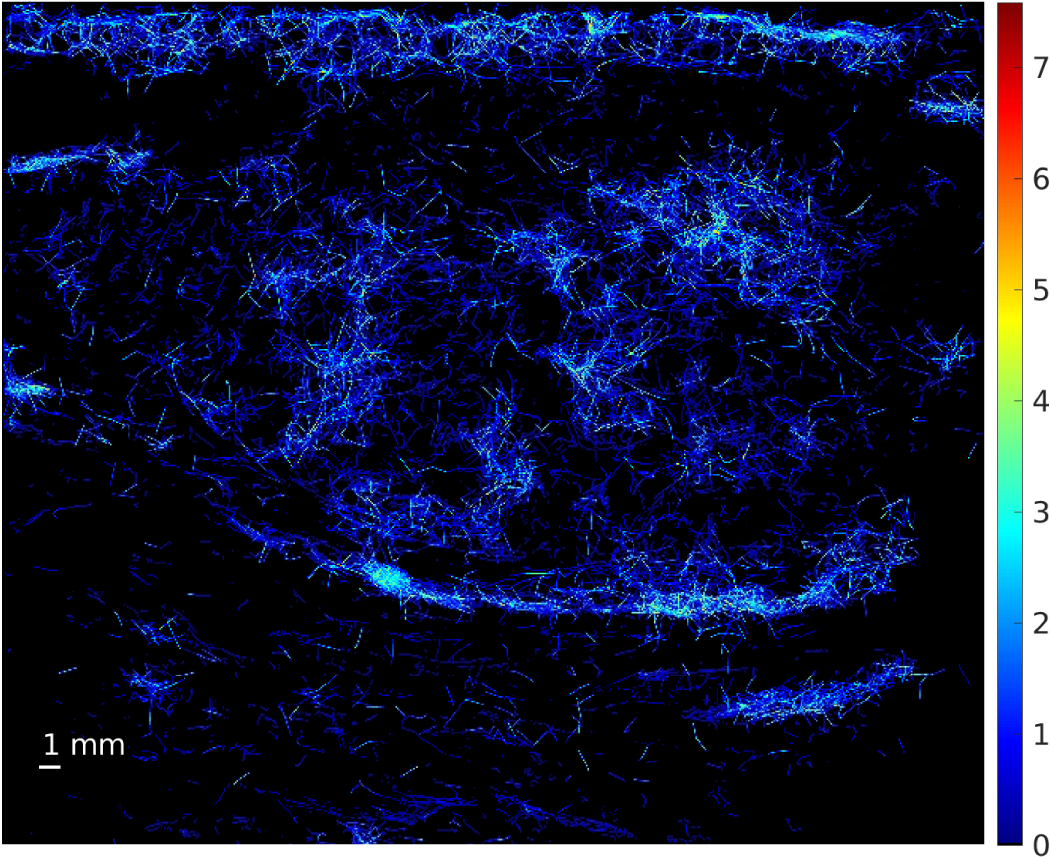
Lymph node MBs velocity magnitude map (mm/s).

### C. Model running time of MBs localization

In this sub-section, a brief comparison between running time of our deep learning based localization and local maxima detection is shown in Table. I to Table. IV. We only report running time per frame in the following tables.

**TABLE I.** Running time of deep learning methods and local maxima detection for synthetic dataset 1

| Method | Image size | Running time (ms) |
| --- | --- | --- |
| nnU-net | $3066 \times 1536$ | 43 |
| ResNet-18 U-net | $3066 \times 1536$ | 31 |
| local maxima detection | $512 \times 512$ | 11 |

**TABLE II.** Running time of deep learning methods and local maxima detection for synthetic dataset 2

| Method | Image size | Running time (ms) |
| --- | --- | --- |
| nnU-net | $1536 \times 1536$ | 9 |
| ResNet-18 U-net | $1536 \times 1536$ | 13 |
| local maxima detection | $512 \times 512$ | 11 |

**TABLE III.** Running time of deep learning methods and local maxima detection for in-vivo rat brain

| Method | Image size | Running time (ms) |
| --- | --- | --- |
| nnU-net | $498 \times 378$ | 0.5 |
| ResNet-18 U-net | $498 \times 378$ | 0.5 |
| local maxima detection | $166 \times 126$ | 0.9 |

**TABLE IV.** Running time of deep learning methods and local maxima detection for in-vivo lymph node

| Method | Image size | Running time (ms) |
| --- | --- | --- |
| nnU-net | $1122 \times 1308$ | 3 |
| ResNet-18 U-net | $1122 \times 1308$ | 3 |
| local maxima detection | $374 \times 436$ | 5 |

Considering that the deep model predictions were performed on 3 times up-sampled images and with sliding windows fashion, the running time for deep model is longer than local maximum detection for some datasets is reasonable. Local maxima detection based localization normally needs extra post-processing of MBs detection for increasing the accuracy of localization, which are normally not necessary for deep learning end-to-end training mode. The extra time for post-processing one frame is different based on the selected methods (see [6] for more details), normally from 3 ms to 48 ms (Gaussian curve fitting) or even more for synthetic dataset. Here we would like to point out that our deep learning based methods were running on GPU (although not so powerful) while the local maxima detection was performed on CPU (Intel Core I7-10700). Thus the comparison may not be rigorously fair.

To the in-vivo rat brain and lymph node set, similar trends as synthetic dataset can be observed. The extra post-processing time per frame for in-vivo rat brain is from around 10 ms to 60 ms. To the in-vivo lymph node dataset, the extra postprocessing time ranges from 18 ms to over 100 ms.

## IV. Discussions and conclusions

In the challenge, the efficiency of our proposed deep learning based MBs localization is preliminarily proved. However, there are still lots of issues that need to be addressed in the future. For example, accurate multi-physics simulations including flow simulation, MBs oscillations induced by ultrasound wave as well as ultrasound beam-forming specialized to SR ultrasound imaging etc. have to be investigated for easily generating abundant training samples. The model architectures need to be further specialized to MBs localization task maybe with the help of Auto ML etc.

We also would like to point out the datasets in the challenge may not well reflect the situations in various clinical or experimental settings of ultrasound imaging. Herein, more data still needs to be acquired and analyzed in order that SR ultrasound imaging can be deployed in our machine.

## Footnotes

[1 Since the challenge focused on the localization and tracking of micro-bubbles in SR ultrasound imaging, below we mainly state the results related with these two parts only. For complete view of SR ultrasound imaging, readers can refer [5] for more details.

